# A direct link between *Prss53*, hair curvature, and skeletal dysplasia

**DOI:** 10.1101/560847

**Authors:** Jichao Deng, Yuning Song, Hongmei Liu, Tingting Sui, Mao Chen, Yuxin Zhang, Bing Yao, Yuxin Xu, Zhiquan Liu, Liangxue Lai, Zhanjun Li

**Author notes:** To whom correspondence should be addressed; (Zhanjun Li); (Liangxue Lai).

## Abstract

In humans, protease serine S1 family member 53 (Prss53) is highly expressed in the hair follicle, especially the inner root sheath, which is associated with hair shape according to recent genome-wide association study (GWAS) data. However, no animal evidence has indicated a link between *Prss53* and hair shape to date. Here, we used CRISPR/Cas9 to generate *Prss53*-mutated rabbits. The homozygous (*Prss53*^-/-^) rabbits exhibited curved hair and skeletal dyskinesia with severe malformation, while the heterozygous (*Prss53*^+/-^) rabbits did not exhibit these features. The curvature features of the hair were accompanied by lesions that were generally denser and less well-defined in the cuticular septation of the hair shaft, and the compartments of the hair follicle were incomplete, as evidenced by decreased expression levels of keratinocyte differentiation genes. In addition, skeletal dysplasia, an increased lethality rate and decreased plasma calcium and serum alkaline phosphatase (ALP) levels were determined in the *Prss53*^-/-^ rabbits. Furthermore, disrupted calcium metabolism, which may play a role in the hair curvature and skeletal dysplasia of *Prss53*^+/-^ rabbits, was demonstrated by using high-throughput RNA sequencing data. Thus, our study confirmed for the first time that the loss of *Prss53* lead to curved hair in animals and provides new insights into the crucial role of *Prss53* in calcium metabolism.

**Author Summary:** No animal evidence has indicated a link between *Prss53* and hair shape to date.

The *Prss53*^-/-^ rabbits exhibited curved hair and skeletal dyskinesia.

The disrupted calcium metabolism may play a role in the hair curvature and skeletal dysplasia of *Prss53*^+/-^ rabbits.

## Introduction

Membrane-anchored serine proteases regulate fundamental cellular and developmental processes, including orchestrating neural tube closure, erecting barriers between cells, developing the inner ear, regulating apical sodium entry and fluid volume, controlling the natriuretic peptide system, adjusting cellular iron export to satisfy iron needs, and promoting fertilization^1^. PRSS53, a member of the membrane-anchored serine proteases, hydrolyzes peptide bonds^2^ and is highly expressed in the hair follicle IRS, medulla, precortex and some melanocytes^3^.

Previous studies reported that *Prss53* is a psoriasis susceptibility locus and is linked to systemic lupus erythematosus, suggesting its potential roles in skin function^4–7^. The Q30R SNP in *Prss53* may influence hair shape and is involved in the human variation of head hair shape among different continental groups^3,8^. In addition, ENaC activity is regulated by the activity of a channel-activating protease of Prss8 (a homologous gene of *Prss53*), implying the *Prss53* may have an indispensable role in ion channel activity^9^.

The triggering of hair curvature is clearly related to symmetry or asymmetry axis formation in follicles and includes Wnt, TGFβ, BMP, Shh and FGFs, etc. ^10,11^. In particular, the Wnt/Ca^2+^ and Wnt/planar cell polarity (PCP) pathways are additional novel mechanisms that affect hair shape^12^. Furthermore, disruptions in calcium metabolism may contribute to a wide variety of clinical symptoms, including osteoporosis, osteolysis, nephrocalcinosis and adynamie^13^.

Here, we successfully generated *Prss53*-deficient rabbits. These rabbits exhibited curved hair, severe skeletal malformation and disrupted calcium metabolism. Altogether, these data suggest that *Prss53* may play an important role in calcium metabolism and provide the first direct evidence that *Prss53* functions in hair curvature shape and skeletal dysplasia.

## Results

### *Prss53* expression pattern

To explore the function of the *Prss53* gene, the expression pattern was determined, as shown in Figure 1. The EST profile data suggest extensive expression of *Prss53* in the skin and bone in Homo sapiens. In addition, high *Prss53* expression in the developing inner root sheath (IRS), precortex and medulla of the hair follicle has been reported in a previous study^3^, but the *Prss53* expression pattern in bone has not been determined.

**Figure. 1.**
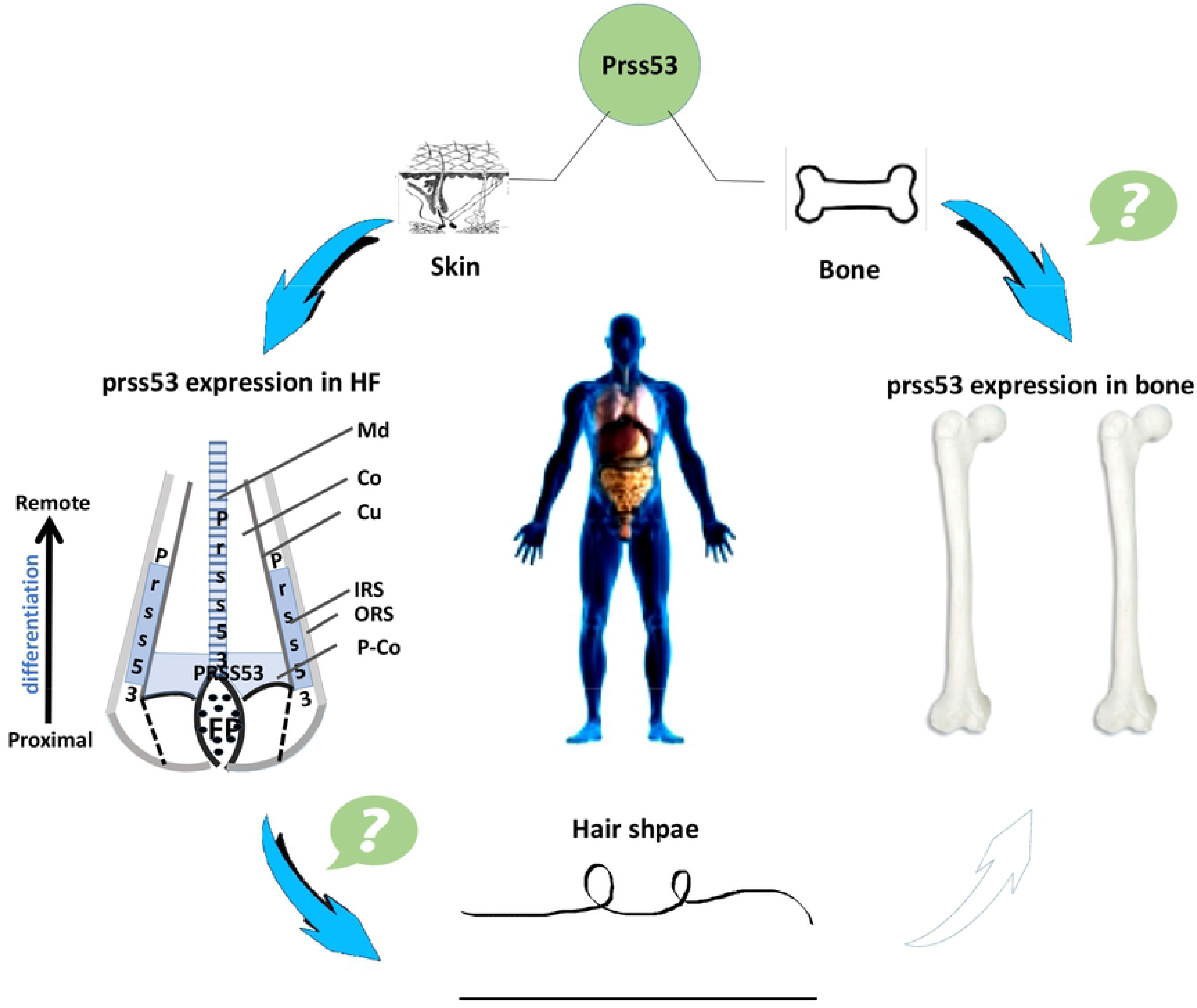
*Prss53* expression pattern. Schematic diagram of the extensive expression of *Prss53* in the lung, stomach, brain, eye, and skin. Bone expression was not determined by the UniGene database. The expression of *Prss53* was mainly detected in the developing IRS, precortex and medulla of the hair follicle.

### Generation of *Prss53*^-/-^ rabbits via the CRISPR/Cas9 system

*Prss53*, a candidate gene for curly hair, was found in a recent GWAS in Latin Americans of mixed European and Native American origin^3^. To disrupt the function of rabbit *Prss53,* two sgRNAs targeting the third and fourth exon of *Prss*53 were designed (Figure 2A) and transferred to embryos. Then, the genomic DNA from each pup was isolated, and *Prss53* gene mutations were detected via PCR and Sanger sequencing. As shown in Figure S1B, F0 rabbits carrying *Prss53* mutations were obtained and were used for the following study. An off-target assay showed that the CRISPR/Cas9 system did not induce undesirable off-target effects in the *Prss53* knockout (KO) rabbits (Supplementary Material, Figure. S2).

**Figure. 2.**
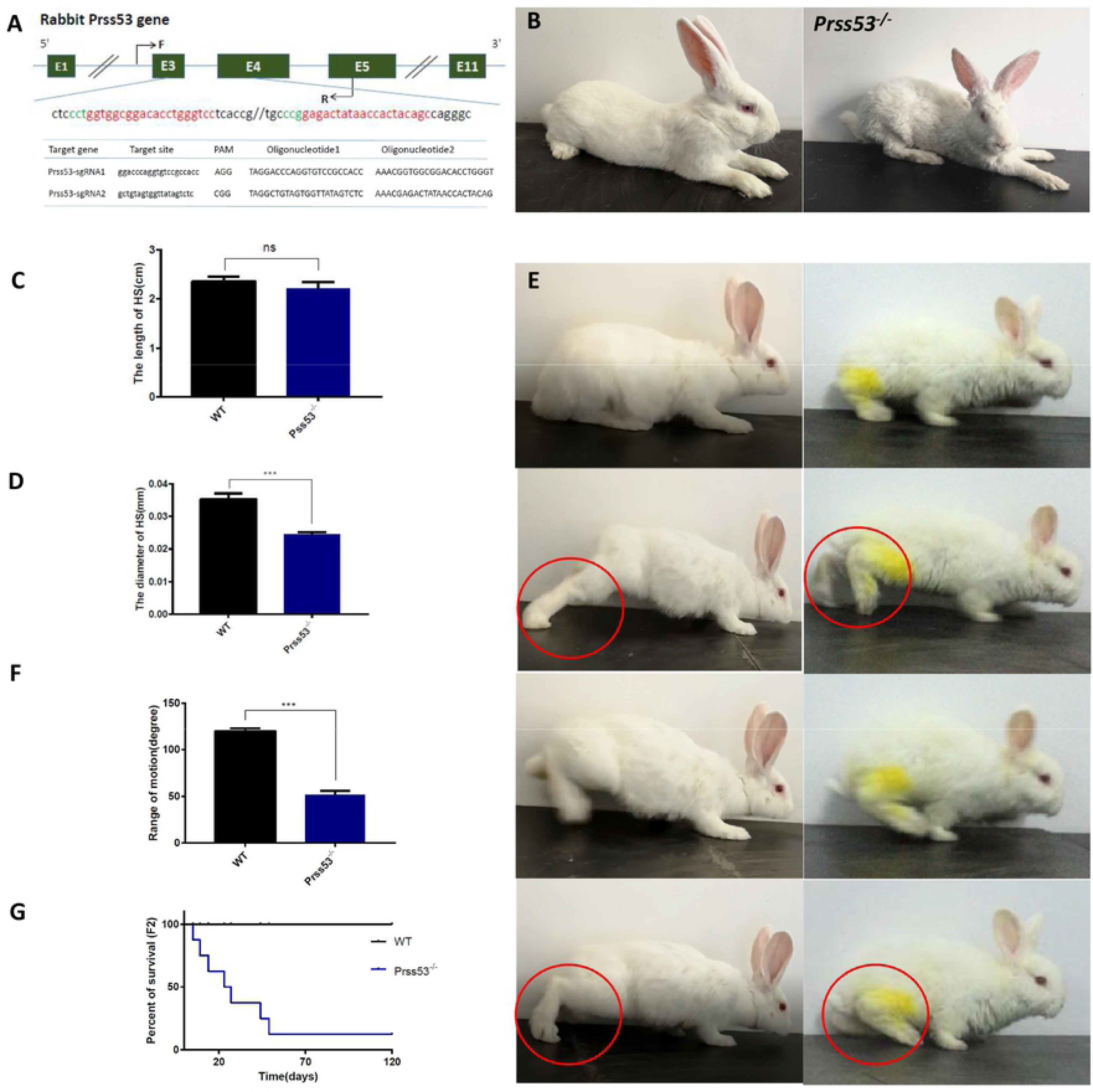
Hair curvature and skeletal abnormalities in *Prss53*^-/-^ rabbits. **(A)** Schematic diagram of two sgRNA target sites located in exons 3 and 4 of rabbit *Prss53*. *Prss53* exons are indicated by rectangles; the target sites of the two sgRNA sequences, namely, sgRNA1 and sgRNA2, are highlighted in red; the protospacer-adjacent motif (PAM) sequence is highlighted in green. **(B)** The hair curvature in a F0 *Prss53*^-/-^ rabbit generated by the CRISPR/Cas9 system. **(C)** No significant difference in the hair length was determined in *Prss53^-/-^* rabbits compared with WT rabbits. **(D)** Statistical analysis showed a decreased hair diameter in the head of *Prss53*^-/-^ rabbits. **(E)** Impaired physical activity was determined in *Prss53^-/-^* rabbits compared with WT rabbits. **(F)** Statistical analysis showed a decreased range of motion in the ankles of *Prss53*^-/-^ rabbits. **(G)** Survival curves of *Prss53*^-/-^ rabbits compared with WT controls.

To obtain a sufficient number of rabbits for a detailed functional study of *Prss53*, the sexually mature F0 founder was mated with wildtype (WT) female rabbits, and then the heterozygous (*Prss53*^+/-^) rabbits were mated and used to generate homozygous (*Prss53*^-/-^) rabbits (Figure. S3). These results demonstrated that mutations in *Prss53* can be achieved via the CRISPR/Cas9 system with high efficiency in rabbits.

### Hair curvature and skeletal dysplasia in *Prss53*^-/-^ rabbits

*Prss53* encodes a polyserine protease called polyserase-3 (POL3S) and is expressed in the IRS, medulla, and precortex of hair follicles and in melanocytes^3^. In our study, curved hair curvature was present all over the body of *Prss53*^-/-^ rabbits (Figure. 2B and S1A). The hair length was not different among the rabbits, but the *Prss53*^-/-^ rabbits had significantly finer hair than that of the WT rabbits (Figure. 2C and D). In addition, impaired physical activity was determined in the *Prss53*^-/-^ rabbits, with a larger <u>knee</u> range of motion compared to the WT rabbits (Figure. 2E and F). The *Prss53*^-/-^ rabbits exhibited a significantly increased lethality rate during postnatal week 6 (~12.5% survive to adulthood) (Figure. 2G).

To further study whether the protein level of *Prss53* was disrupted in the *Prss53*^-/-^ rabbits, we performed a 3D model protein structure prediction, Western blotting and immunofluorescence staining. The results indicated a disrupted protein structure (Figure. 3A and B). The complete loss of PRSS53 protein (Figure. 3C and D) was determined in the *Prss53*^-/-^ rabbits when compared with the WT rabbits. Furthermore, the immunohistochemical results confirmed the complete loss of PRSS53 protein in the hair follicles (Figure. 3E and F) and bone marrow (Figure. 3G and H) of the *Prss53*^-/-^ rabbits. These results confirmed that *Prss53* plays an important role in maintaining hair shape and skeletal health.

**Figure. 3.**
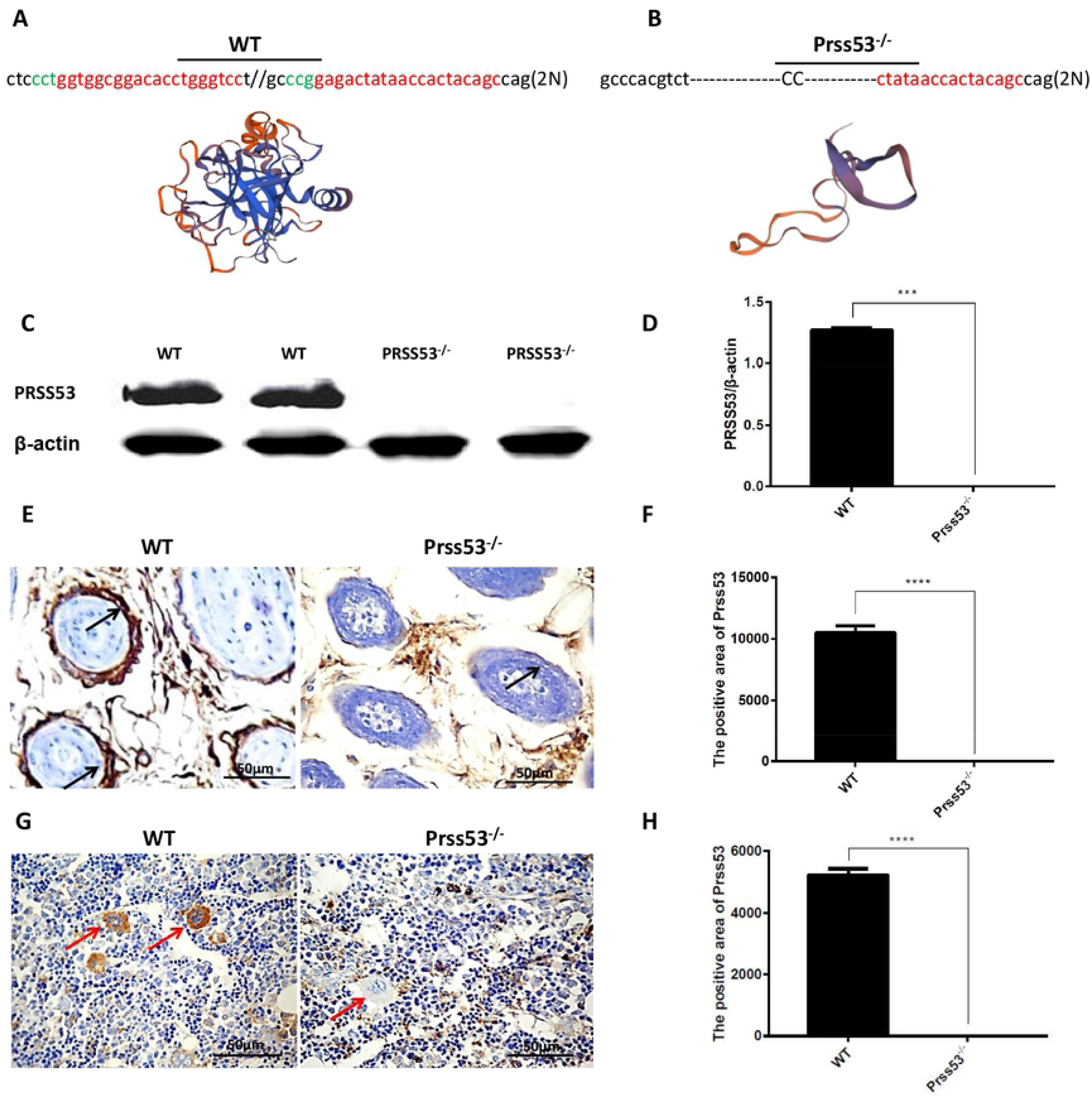
The loss of PRSS53 protein in *Prss53*^-/-^ rabbits. **(A and B)** Computer modeling of the 3D structure of WT and mutant *Prss53*. **(C and D)** Western blot results showed the complete loss of PRSS53 protein in the skin of *Prss53*^-/-^ rabbits. **(E and F)** Immunohistochemistry staining showed the complete loss of PRSS53 protein in the hair follicles of *Prss53*^-/-^ rabbits. Black arrow indicates the expression of PRSS53 protein in the hair follicles of WT rabbits. **(G and H)** Immunohistochemistry staining showed the complete loss of PRSS53 protein in the proerythroblasts of *Prss53*^-/-^ rabbits. Red arrow indicates the expression of PRSS53 protein in the proerythroblasts of WT rabbits.

### Abnormal keratinocyte differentiation in *Prss53*^-/-^ rabbits

A previous study reported that abnormal keratinocyte differentiation negatively affects the hair shaft extension and hair shaft shape^14^. In this study, H&E staining showed no significant pathological changes of the hair follicles in the *Prss53*^-/-^ rabbits (Figure. 4A). Next, we examined the hair structure by scanning electron microscopy (SEM). As illustrated in Figure. 4C, the medullated fibers and unmyelinated hair from WT rabbits exhibited well-defined and regular cuticular septation, while the *Prss53*^-/-^ rabbits presented lesions and less well-defined cuticular septation. Furthermore, transmission electron microscopy (TEM) analyses were performed on the skin from the dorsum of *Prss53*^-/-^ and WT rabbits. The results showed that the WT rabbits exhibited a clear structure, including IRS and cuticle, while the *Prss53*^-/-^ rabbits fail to develop the cuticle (Figure. 4D). High-throughput sequencing revealed significantly decreased expression of keratinocyte differentiation genes, including *Lats1, Yap1, Adam9, Csta, Krt10, Wnt16 and Tp63,* in the *Prss53*^-/-^ rabbits compared with the WT rabbits (Figure. 4B). The results showed abnormal keratinocyte differentiation in the *Prss53*^-/-^ rabbits.

**Figure. 4.**
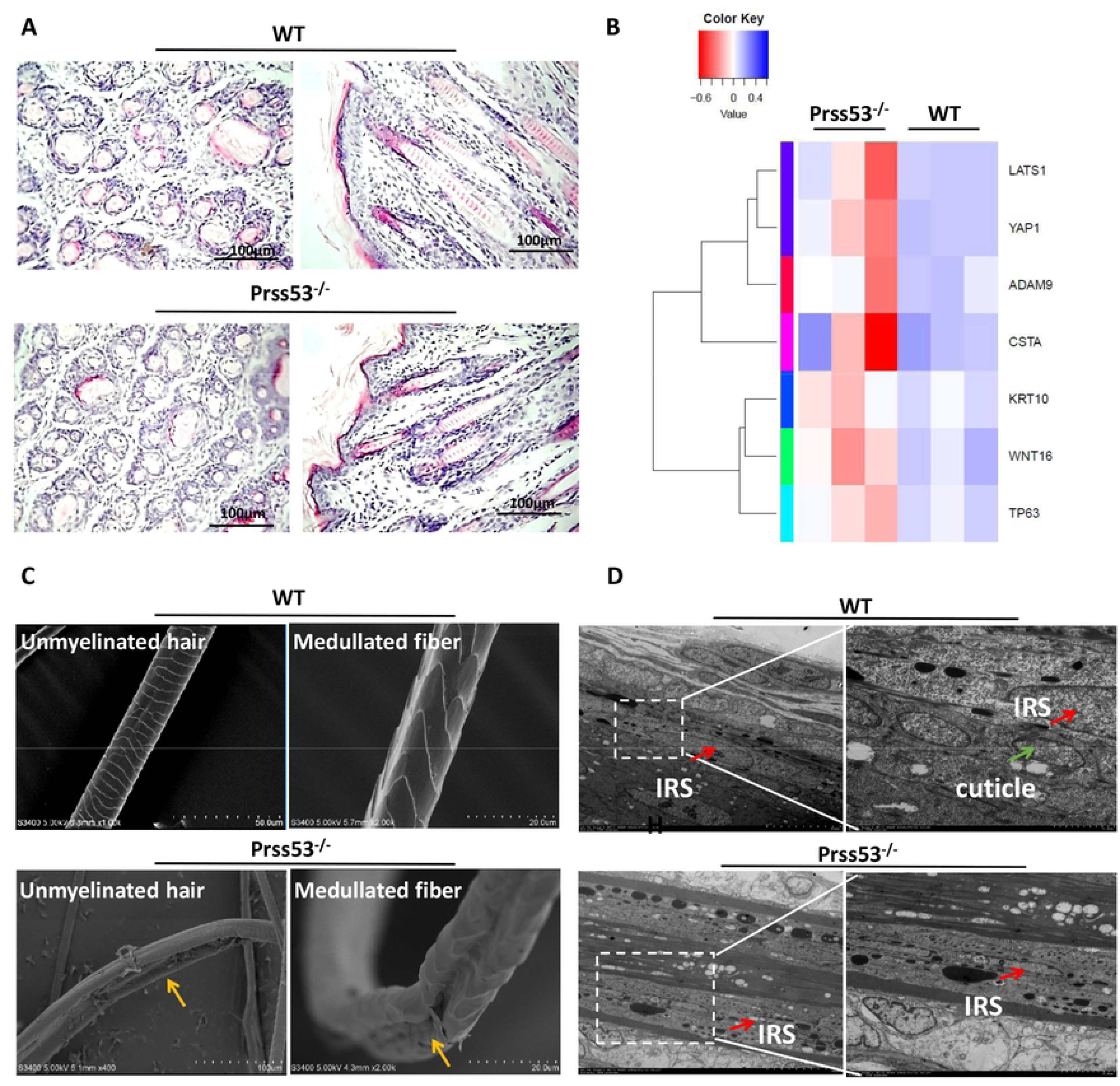
Abnormal keratinocyte differentiation in *Prss53^-/-^* rabbits. (A) No significant difference in the HF was determined in the *Prss53*^-/-^ rabbits compared with WT rabbits. **(B)** The heat map shows the significantly decreased expression of keratinocyte differentiation genes, including LATS1, YAP1, ADAM9, CSTA, KRT10, WNT16 and TP63, in *Prss53*^-/-^ rabbits compared with WT rabbits **(C)** SEM analysis of WT and *Prss53*^-/-^ hair fibers shows that medullated fibers and unmyelinated hair from WT rabbits exhibit well-defined and regular cuticular septation, while medullated fibers and unmyelinated hair from *Prss53*^-/-^ rabbits reflect lesions that appear generally denser and less well-defined areas in the cuticular septation. **(D)** TEM analysis of WT and *Prss53*^-/-^ skin shows that WT rabbits exhibited a clear structure, while the *Prss53*^-/-^ rabbits fail to develop the cuticle. Red arrow indicates the location of the IRS in hair follicle of WT and *Prss53*^-/-^ rabbits; green arrow indicates the location of the cuticle in the hair follicle of WT rabbits; yellow arrow indicates lesions and less well-defined areas in the cuticular septation.

### Disrupted calcium metabolism in *Prss53*^-/-^ rabbits

Extracellular calcium and 1,25(OH)_2_D raise intracellular free calcium as a necessary step toward stimulating keratinocyte differentiation^15^. To uncover whether *Prss53* influences hair shape by regulating calcium metabolism, high-throughput RNA sequencing was used to compare the gene expression patterns between the *Prss53*^-/-^ and WT rabbits. As shown in Figure. 5A, good internal controls and high correlations were determined in the *Prss53*^-/-^ and WT rabbits. The up-regulated and down-regulated genes are shown in Figure. 5B. Furthermore, Gene Ontology (GO) and Kyoto Encyclopedia of Genes and Genomes (KEGG) analyses revealed significantly upregulated calcium ion binding genes and the calcium signaling pathway in the *Prss53*^-/-^ rabbits compared with the WT rabbits (Figure. 5C and Table 1). Thus, we suspected that abnormal calcium metabolism is responsible for the hair curvature phenotype of the *Prss53*^-/-^ rabbits (Figure. 5D).

**Table 1:**
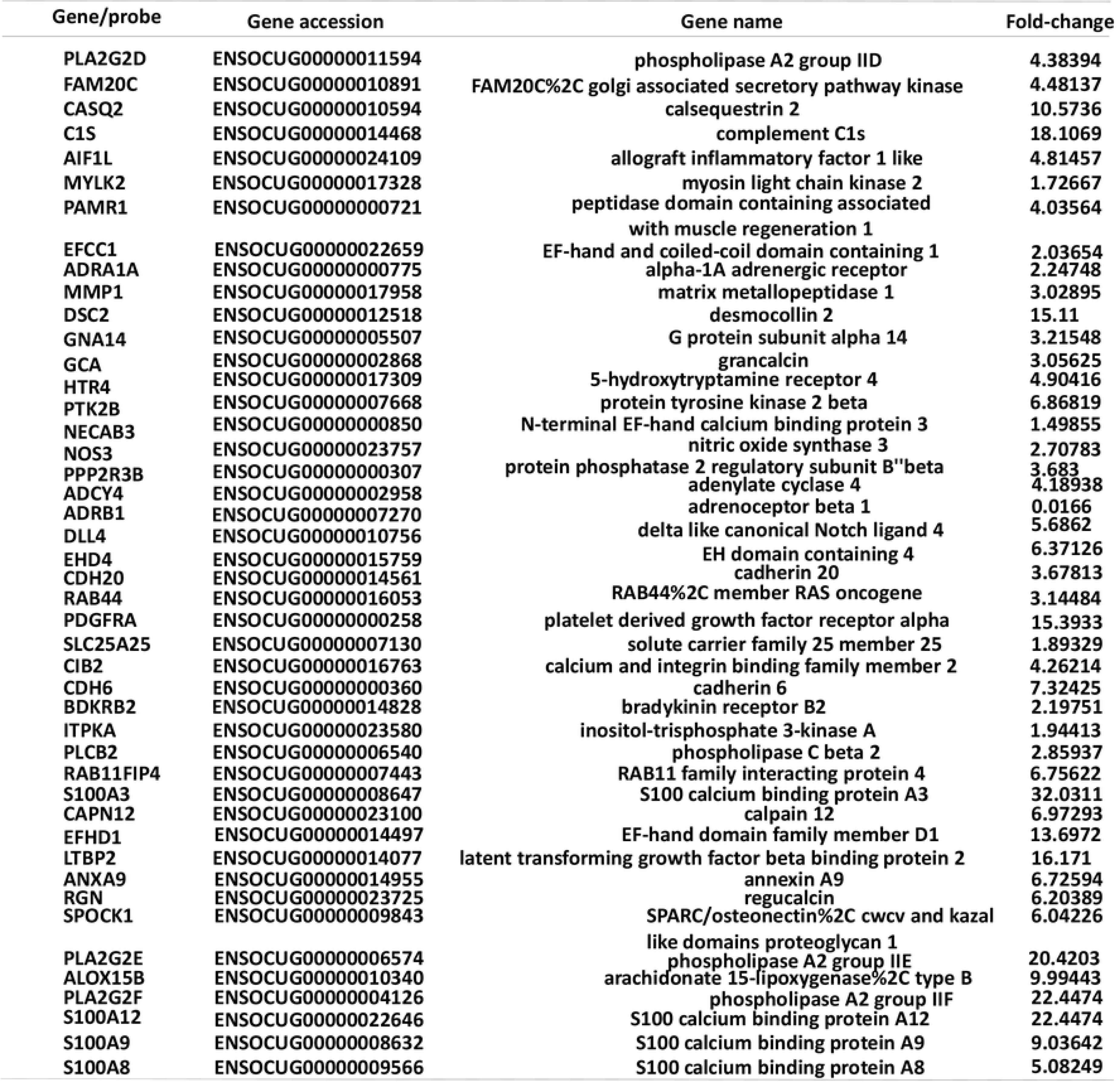
The expression of calcium metabolism gene.

**Figure. 5.**
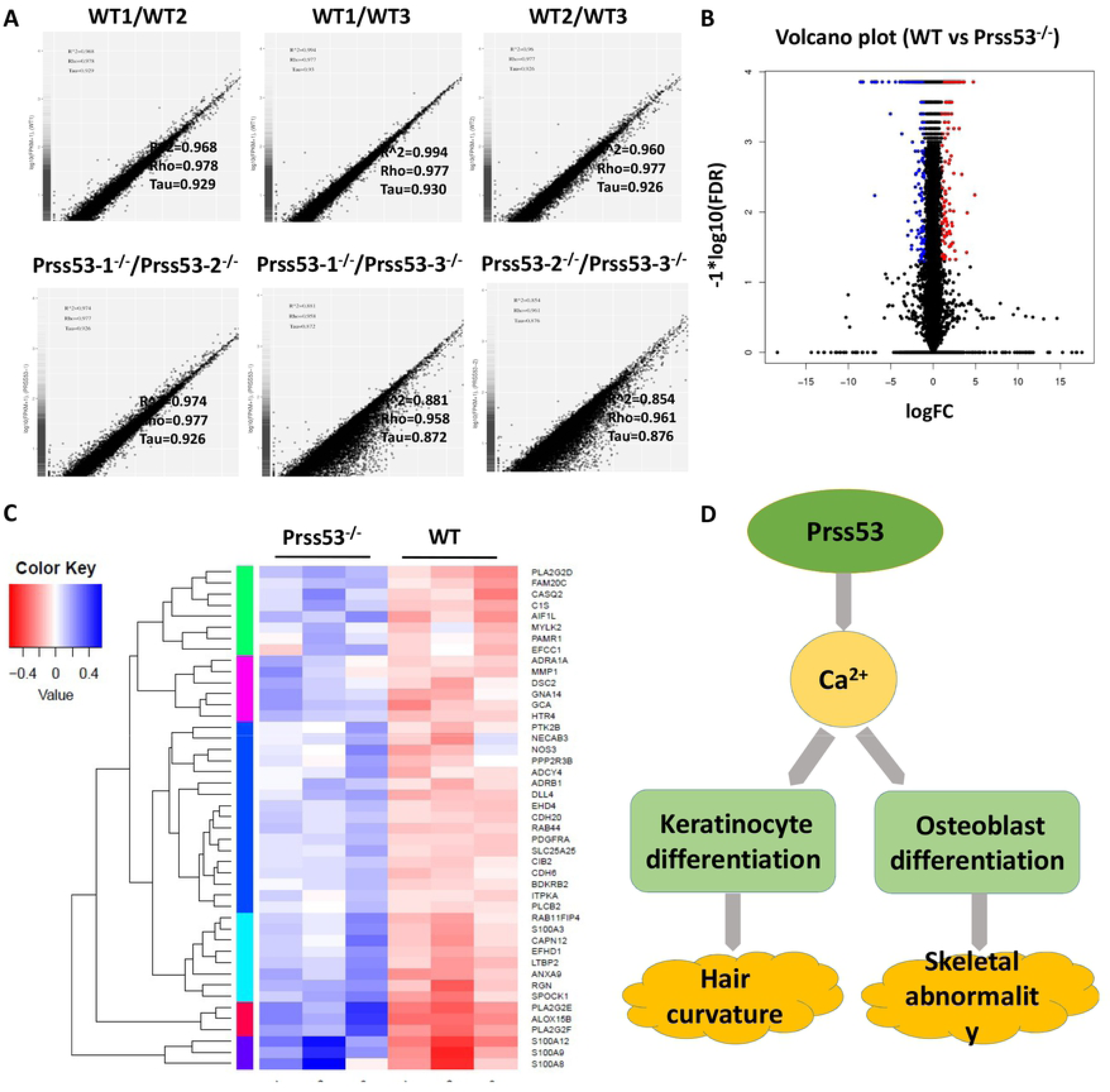
Disrupted calcium metabolism in *Prss53*^-/-^ rabbits. (A) Internal consistency, homogeneity and high correlations were determined in the WT and *Prss53*^-/-^ rabbits. **(B)** Volcano plot shows that a large number of differentiated genes (upregulation and downregulation) was detected between WT and *Prss53*^-/-^ rabbits. Blue dots indicate upregulated genes; red dots indicate downregulated genes. **(C)** Heap map showing significantly upregulated calcium metabolism genes in the *Prss53*^-/-^ rabbits compared with WT rabbits. **(D)** Schematic diagram of the role of *Prss53* in calcium metabolism, which is responsible for keratinocyte differentiation and osteoblast differentiation to maintain the normal morphology of the hair and skeleton.

### Disrupted osteoblast differentiation in *Prss53*^-/-^ rabbits

Disordered calcium metabolism may induce osteoporosis and osteolysis in clinical settings^13^. To further examine whether disrupted calcium metabolism is responsible for the skeletal dysplasia, the skeletal and a serum biochemical analysis were compared between the *Prss53*^-/-^ and WT rabbits. As shown in Figure. 6A, the *Prss53*^-/-^ rabbits exhibited more severe knee abnormalities and were substantially more malformed than WT rabbits by X-ray autoradiograph examination. The serum biochemical analysis demonstrated significantly decreased plasma calcium and ALP levels in the *Prss53*^-/-^ rabbits (Figure. 6B and C). In addition, immunohistochemistry and H&E staining confirmed the significantly reduced number of osteoblasts and the complete loss of PRSS53 protein in the *Prss53*^-/-^ rabbits (Figure. 6D, E and F) compared to the WT control. Hence, disrupted calcium metabolism is responsible for the skeletal abnormalities in the *Prss53*^-/-^ rabbits.

**Figure. 6.**
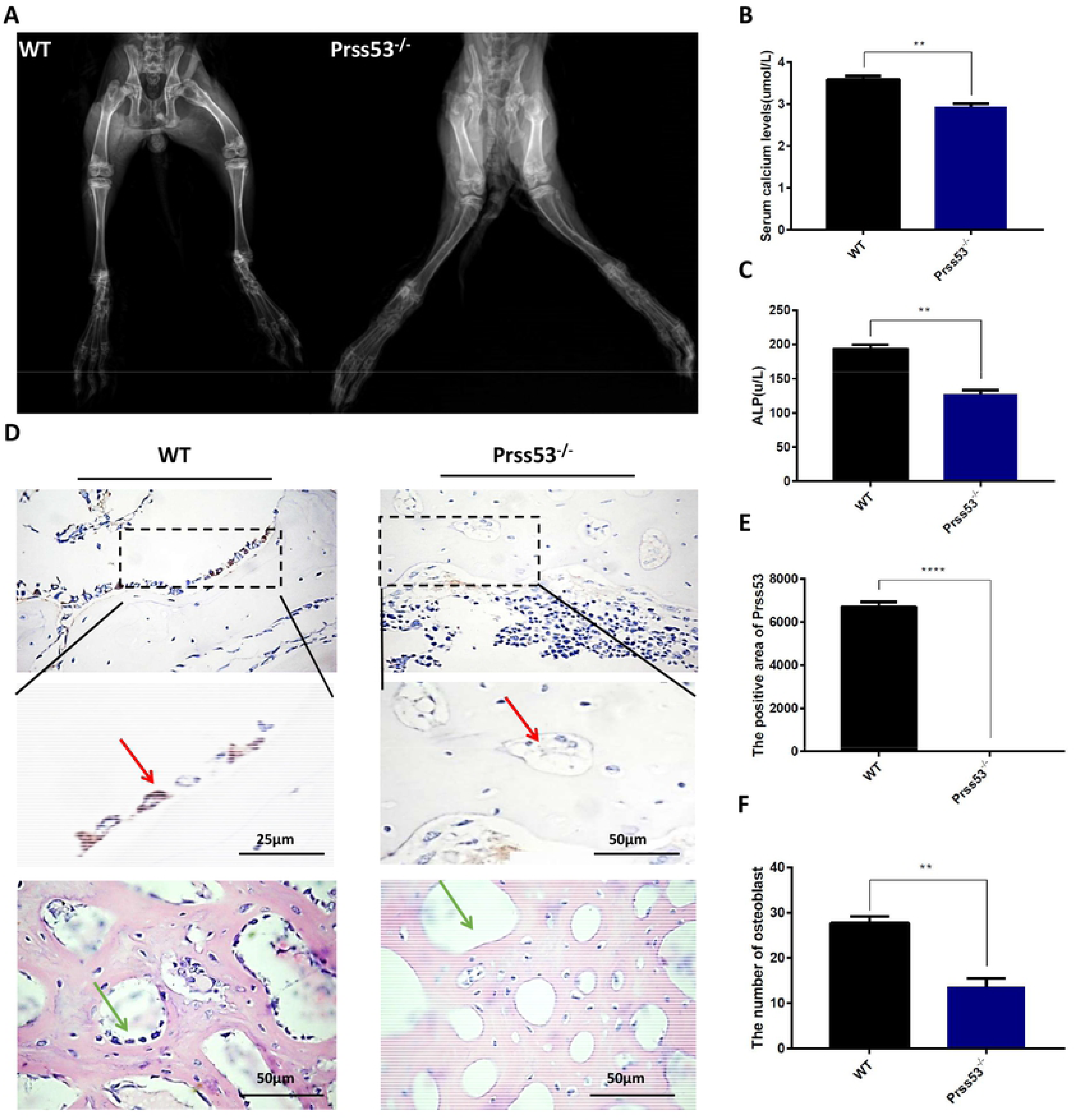
Disrupted osteoblast differentiation in *Prss53*^-/-^ rabbits. **(A)** X-ray autoradiography was used to characterize the skeleton in WT and *Prss53*^-/-^ rabbits. **(B and C)** Decreased bone remodeling markers of serum Ca^2+^ and serum alkaline phosphatase (ALP) in the *Prss53*^-/-^ rabbits. **(D)** H&E and immunohistochemistry staining showed the high expression of *Prss53* in the osteoblasts, although not in the *Prss53*^-/-^ rabbits. A significantly reduced number of osteoblasts was determined in the *Prss53*^-/-^ rabbits by H&E staining. **(E)** Statistical analysis showing PRSS53 expression in the osteoblasts, although not in the *Prss53*^-/-^ rabbits. **(F)** Statistical analysis showing a significantly reduced number of osteoblasts in the *Prss53*^-/-^ rabbits. Red arrow indicates the expression of PRSS53 protein in the osteoblasts of WT rabbits. Green arrow indicates the location of osteoblasts in WT rabbits.

In summary, our study provides the first evidence that *Prss53* plays an important role in calcium metabolism, which is responsible for keratinocyte and osteoblast differentiation and, thereby, maintains the normal morphology of the hair and skeleton (Figure. 5D).

## Discussion

To date, the function *Prss53* was largely unknown, and limited information regarding *Prss53* mutant animals was available. In this study, we first generated a stably transmitted *Prss53* mutant rabbit via the CRISPR/Cas9 system. Interestingly, the hair curvature phenotype of the *Prss53*^-/-^ rabbits is consistent with the Latin American population of mixed European and Native American origin with a Q30R substitution in *Prss53* ^3^.

Uncombable hair syndrome is characterized by dry, frizzy, and spangly hair, which has longitudinal grooves and a heart-shaped cross section by SEM^16^. Consistently, lesions and less well-defined cuticular septation were determined in the *Prss53*^-/-^ rabbits in our study. In addition, high-throughput RNA sequencing data revealed significantly decreased expression of keratinocyte differentiation genes, including *Lats1*^17^, *Yap1*^18^, *Adam9*^19^, *Csta*^20^, *Krt10*^21^, *Wnt16*^22^ and *Tp63*^23^, in *Prss53*^-/-^ rabbits compared with WT rabbits, A previous study showed that the hair shaft is derived from the progeny of keratinocyte stem cells in the follicular epithelium, and the growth and differentiation of follicular keratinocytes is guided by a specialized mesenchymal population^14^. Therefore, abnormal keratinocyte differentiation is responsible for disruptions to hair shaft shape, indicating that *Prss53* may play an important role in maintaining keratinocyte differentiation.

Unexpectedly, impaired physical activity and decreased plasma calcium may be responsible for the early death of *Prss53*^-/-^ rabbits. Calcium is essential for bone development by promoting osteogenesis and inhibiting osteoclast activity through increased osteoprotegerin secretion from osteoblasts^24^. Alkaline phosphatase is widely used as a marker of osteoblast differentiation and was significant decreased in the *Prss53*^-/-^ rabbits when compared with WT rabbits. Previous studies reported that osteoblast differentiation is closely associated with skeletal development, such as in *Med23-*deficient MSCs or preosteoblasts that display defective bone formation and impaired osteoblast differentiation^25^. *Runx2*^−/−^ mice lack both mature osteoblasts and a mineralized skeleton^26,27^. Hence, this study is the first report to demonstrate that disrupted osteoblast differentiation in the *Prss53* mutant is responsible for skeletal abnormalities, suggesting that *Prss53* may have an indispensable role in skeletal development.

In this study, the significantly reduced number of osteoblasts and disrupted calcium metabolism were determined in the *Prss53*^-/-^ rabbits by RNA sequencing technology. The G-protein-coupled receptor (CaR) is not limited to the parathyroid gland ^28^, and also inkeratinocytes^29^. The calcium-sensing receptor, transient receptor potential channels, and STIM/Orai also function in calcium sensing and calcium entry in keratinocytes^30^. In addition, both extracellular calcium and 1,25(OH)_2_D raise the level of intracellular free calcium as a necessary step toward stimulating keratinocyte differentiation^15^. Accumulating studies have shown that calcium oscillations via restraints of major Ca^2+^ entry sources (extracellular Ca^2+^ influx and intracellular Ca^2+^ release from the endoplasmic reticulum) is required for osteoblast differentiation^31^. Hence, we hypothesize that disrupted calcium metabolism is responsible for abnormal keratinocyte differentiation in the *Prss53*^-/-^ rabbits.

To the best of our knowledge, this study is the first animal report on hair curvature and skeletal dysplasia caused by a *Prss53* mutation in rabbits and provides evidence that *Prss53* is required for keratinocyte and osteoblast differentiation, which is essential for hair and skeletal cell proliferation.

## Materials and Methods

### 1 Ethical statement

New Zealand rabbits were obtained from the Laboratory Animal Center of Jilin University. All rabbit experiments were conducted under the approval of the Animal Care Center and Use Committee of Jilin University.

### 2 sgRNA design and vector construction

The 3xFLAG-NLS-SpCas9-NLS vector (Addgene ID 48137) was linearized with *NotI* and *in vitro* transcribed using a mMessage mMachine SP6 Kit (Ambion) and RNeasy Mini Kit (Qiagen), according to the manufacturer’s instructions.

The design of targeted sgRNAs was described previously (http://crispr.mit.edu/)^32^. Complementary oligo sgRNAs were cloned into the *BbsI* sites of a Puc57-T7-sgRNA cloning vector (Addgene ID 51306). The Puc57-T7-sgRNA vector was amplified by PCR with T7 primers (T7-F: 5’-GAA ATT AAT ACG ACT CAC TAT A-3’ and T7-R: 5’-AAA AAA AGC ACC GAC TCG GTG CCA C-3’), and then, the PCR products were *in vitro* transcribed using a MAXIscript T7 Kit (Ambion, USA) and purified with a miRNeasy Mini Kit (Qiagen, Germany), according to the manufacturer’s instructions.

### 3 Microinjection and embryo transfer

The procedures of embryo microinjection and embryo transfer were previously described ^33,34^. Briefly, female New Zealand white rabbits (6–8 months old) were superovulated with FSH (50 IU) every 12 h for 3 days. After the last injection, the female rabbits were mated with male rabbits and were then injected with 100 IU of human chorionic gonadotrophin (hCG). Rabbit embryos at the pronuclear stage were collected and transferred into oocyte manipulation medium. A mixture of Cas9 and sgRNA mRNA (200 ng/μl and 40 ng/μl, respectively) was microinjected into the embryo cytoplasm. The injected embryos were transferred to EBSS medium for short-term culture at 38.5°C, 5% CO_2_ and 100% humidity conditions. Approximately 30–50 injected zygotes were transferred into the oviducts of recipient rabbits.

### 4 Mutation detection in pups by PCR

Genomic DNA from WT and *Prss53* mutant rabbits was isolated using a TIANamp Genomic DNA Kit (TIANGEN, Beijing, China). The DNA was amplified by 2×Taq Plus MasterMix (TIANGEN), and the PCR primers used to detect mutations were as follows: rabbits-F-5’ CAG GAA GTT CCA GTC ACT TGT −3’, rabbits-R-5’ GGT TGA GAA GGA AGG GAG ATT AG-3’. The PCR products were purified and cloned into the pGM-T vector (TIANGEN, Beijing, China); at least 10 positive plasmid clones were sequenced and analyzed using NCBI BLAST.

### 5 Western blot

Western blot analysis was performed as described previously^35^. Samples from the head skin of *Prss53*^-/-^ and WT rabbits were homogenized and lysed in RIPA buffer supplemented with 2.5 μL/mL protease inhibitor cocktail (Roche) on ice for 30 min. Anti-*Prss53* (1:1000, Novus) were used as primary antibody, and β-actin antibody (1:2000, Abcam) was used as loading control.

### 6 Histopathology and immunohistochemistry

H&E staining was performed as previously described ^33^. Briefly, the tissues were collected from WT and *Prss53^-/-^* rabbits, fixed in 4% paraformaldehyde for 48 h, embedded in paraffin wax, sectioned for slides, and stained with H&E.

The immunohistochemistry staining was performed as previously described^36^. Primary antibodies against *Prss53* (1:500, Novus) and secondary anti-rabbit polyclonal antibodies (1:1000, Beyotime Institute of Biotechnology, Shanghai, China) were used. The slides were imaged with a Nikon ts100 microscope and processed using Photoshop CS5 (Adobe).

### 7 Scanning electron microscopy (SEM)

Scanning electron microscopy (SEM) was performed as described previously^37^. The hair from the heads of *Prss53*^-/-^ and WT rabbits was attached onto specimen stubs using double-sided conductive tabs and sputter-coated with gold using a Polaron SEM E-1010. Samples were imaged using a S-3400N Scanning Electron Microscope.

### 8 Transmission electron microscopy (TEM)

Transmission electron microscopy was performed as described previously^38^. The skin from the dorsum of *Prss53*^-/-^ and WT rabbits was fixed in 3% glutaraldehyde for 4 h at 4°C. Ultrathin longitudinal sections of the skin were cut with an ultramicrotome and a diamond knife and processed for examination by transmission electron microscope (TEM) (H-7640, Hitachi, Japan).

### 9 Skeletal histomorphology

X-ray autoradiography pictures of the *Prss53*^-/-^ and WT rabbits were taken as previously described ^33^. Briefly, the YEMA Radiography System with a digital camera attached (Varian, USA) on X-ray film (ROTANODE, Japan) was used in this study. The images were taken at 40 KV with 3 mA exposure.

### 10 Statistical analysis

Data were statistically analyzed with GraphPad prism software (t test), and *p* < 0.05 was considered statistically significant. \**p* < 0.05, \*\**p* < 0.01, \*\*\**p* < 0.001, **** *p* < 0.0001.

## Abbreviation

Prss53: serine protease 53
ALP: alkaline phosphatase
POL3S: polyserine protease called polyserase-3
CAP: channel-activating protease
PCP: planar cell polarity
IRS: inner root sheath
KO: knockout
WT: wildtype
HF: hair follicle
SEM: scanning electron microscope
TEM: transmission electron microscopy
GO: gene ontology
KEGG: Kyoto encyclopedia of genes and genomes
CaR: G-protein-coupled receptor.

## Acknowledgments

The authors express their gratitude to Peiran Hu at the Embryo Engineering Center for technical assistance.

This study was financially supported by the National Key Research and Development Program of China Stem Cell and Translational Research (2017YFA0105101), the Program for Changjiang Scholars and Innovative Research Team in University (No. IRT_16R32), and Key Research & Development Program of Guangzhou Regenerative Medicine and Health Guangdong Laboratory (2018GZR110104004).

## Author Contributions

J.D., L.L., and Z.L. designed the research. J.D., Y.S., and T.S. performed the research. J.D., H.L., M.C., and B.Y. contributed new reagents or analytic tools. J.D., Y.Z., Y.X., and Z.L. analyzed the data. J.D., Z.L., and L.L. wrote the paper. All authors have read and approved the final manuscript.

**Competing financial interests:** The authors declare no competing financial interests.

**Figure. S1. Generation of *Prss53*^-/-^ rabbits via the CRISPR/Cas9 system (A)** Photographs showing the curved hair of *Prss53*^-/-^ rabbits. **(B)** Agarose gel electrophoresis for mutation detection in F0 generation pups. M: DL2000. Sanger sequencing of the *Prss53* mutant rabbits. The sgRNA sequences are highlighted in red, and PAM sequences are indicated in green. Deletion, “−”.

**Figure. S2. Off target determination of the *Prss53* mutant rabbits** T7E1 cleavage analysis of POTS for sgRNA1 and sgRNA2. M, DL2000; OT11-15 and OT21-25 represent the ten POTS. The chromatogram sequence analysis of ten POTS for sgRNA and sgRNA2 using PCR products in founder rabbits. The 20 bp of the POTS and PAM are represented in the shaded area.

**Figure. S3. Heritability of the *Prss53* mutant rabbits** (A) A male chimera rabbit was mated with female WT rabbits, and there was no hair curvature phenotype in the F1 heterozygous (*Prss53*^+/-^) rabbits. **(B)** The heterozygous (*Prss53*^+/-^) rabbits were mated and used for the generation of homozygous (*Prss53*^-/-^) rabbits. **(C)** Agarose gel electrophoresis for mutation detection of the F1 *Prss53* mutant rabbits. The sgRNA sequences are highlighted in red, and PAM sequences are indicated in green. Deletion, “−”. **(D)** Agarose gel electrophoresis for mutation detection of the F2 *Prss53* mutant rabbits. The sgRNA sequences are highlighted in red, and PAM sequences are indicated in green. Deletion, “−”.

**Table S1: Primers used for qRT-PCR**

**Table S2: Primers used for off target**

